# *WormSNAP*: A software for fast, accurate, and unbiased detection of fluorescent puncta in *C. elegans*

**DOI:** 10.1101/2025.01.28.635348

**Authors:** Araven Tiroumalechetty, Elisa B. Frankel, Peri T. Kurshan

## Abstract

The detection and characterization of fluorescent puncta are critical tasks in image analysis pipelines for fluorescence imaging. Existing methods for quantitative characterization of such puncta often suffer from biases and limitations, compromising the reliability and reproducibility of results. Moreover, the widespread adoption of many available analysis scripts is often hampered by over-optimization for specific samples, requiring extensive coding knowledge to repurpose for other datasets. We present *WormSNAP (Worm SyNapse Analysis Program)*, a freely available, stand-alone, no-code approach to automated unbiased detection and characterization of 2D fluorescent puncta, originally developed to characterize images of the synapses residing in *C. elegans* nerve cords but suitable for broader 2D fluorescence image analysis. *WormSNAP* incorporates a local means thresholding algorithm and a user-friendly Graphical User Interface (GUI) for efficient and accurate analysis of large datasets, with user control of thresholding and restriction parameters and visualization options for further refinement. *WormSNAP* also calculates three types of correlation metrics for 2-channel images, enabling users to select the ideal metric for their dataset. *WormSNAP* provides robust and accurate fluorescent puncta detection in a variety of conditions, accelerating the image analysis workflow from data acquisition to figure generation.

## INTRODUCTION

### Fluorescent puncta detection in *C. elegans* research

*C. elegans* research has progressed in tandem with live fluorescent imaging, beginning with the validation of GFP as a marker for gene expression in eukaryotes^1^. Since then, the genetic tractability of *C. elegans* has enabled the generation of numerous cell-specific fluorescent protein (FP) expression systems, ranging from overexpressed plasmid-based arrays to endogenous CRISPR-based reporters, leading to major discoveries about the molecular basis underlying diverse cellular processes^2^. For example, fluorescence imaging of the *C. elegans* nervous system has yielded significant insights into the roles of synaptic proteins and enabled the dissection of complex cell-biological signaling pathways involved in synapse formation, patterning, maintenance, and plasticity ^3–7^.

Although genetic and microscopy innovations have gradually reduced the barriers to fluorescence image acquisition in the *C. elegans* nervous system, the quantification of fluorescence imaging data remains non-standardized. Segmentation methods for defining fluorescent regions of interest (ROIs) vary widely, and most researchers carry out fluorescent puncta detection manually or through custom scripts, often optimized for a specific reporter, cell morphology, phenotype of interest, or dataset^8,9^. Such strategies can expose findings to experimenter bias and jeopardize reproducibility, as observed in other image analysis contexts^10^. Substantial variation also exists in how commonly reported fluorescence puncta characteristics – such as intensity, distribution, and circularity – are measured and analyzed. Furthermore, most freely available analysis software for fluorescence puncta quantification requires some coding expertise to implement or modify and is infrequently updated or supported, leaving researchers to choose between subpar automated methods or tedious manual annotation.

*C. elegans* exhibit a highly stereotypical body plan and nervous system arrangement (schematized in Figure 1A). Most images of synaptic puncta in *C. elegans* are collected as either widefield fluorescent images or z-stack projections from a confocal microscope. Images of *en passant* synapses of the nerve cords are generally cropped and often straightened with the use of a segmented line tool (producing uniform width, cropped images referred to as *crops* in this paper). A 1D sum projection of each *crop* is then used to generate an intensity plot profile for further analysis (Figure 1Dii) ^9,11,12^. However, reducing images to 1D projections discards much of the spatial information about the fluorescent puncta, restricting the output information primarily to intensity along the process. Although some progress has been made in switching to automated identification of 2D regions of interest (ROIs) to retain spatial information ^8,12^, no widely adopted method yet exists in this subfield^8,12^. Meanwhile, CRISPR-mediated endogenous protein labeling – while less prone to overexpression artifacts – often results in dimmer signals and, consequently, lower signal-to-noise ratios (Figure 1 B, C). This further reduces the suitability of intensity profiles for puncta detection and characterization.

**Figure 1.**
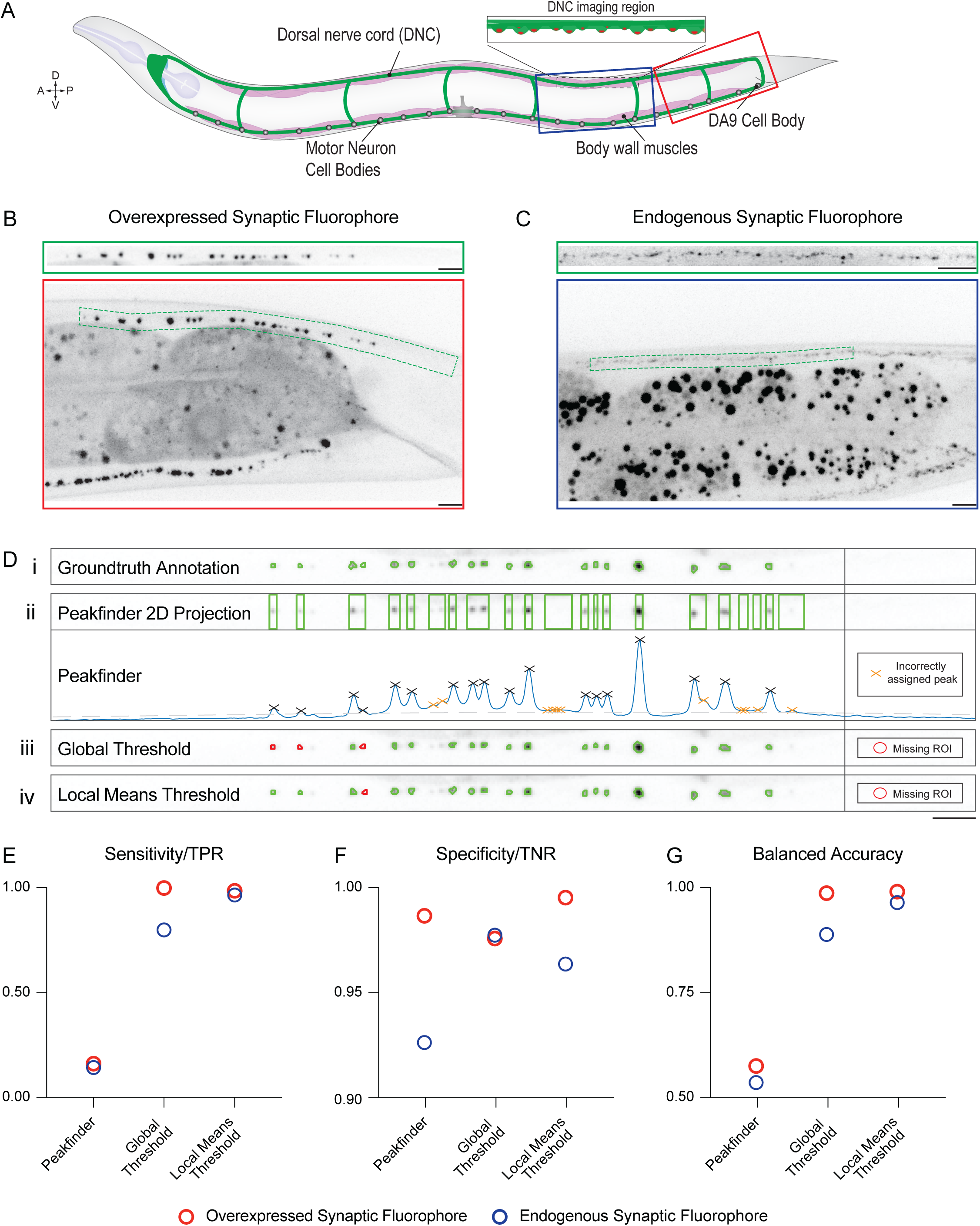
Local Means Threshold shows higher accuracy than current standard ROI detection methods in crops. A. Schematic of *C. elegans* body plan, with imaging regions displayed in B and C boxed in dark blue and red, respectively. B. (Top) Straightened *crop* extracted from the full-size confocal image below. (Bottom) 63X confocal image of an L4-stage worm that is overexpressing the synaptic active zone marker CLA-1::GFP in the cholinergic motor neuron DA9, whose axon extends in the dorsal nerve cord. The crop outline is shown in green. Note the nearby dark signal from the gut (dark gray), including gut granules (dark spots). Scale Bars, 5µm. C. (Top) Straightened *crop* extracted from the same region as displayed in the full-size confocal image below. (Bottom) 63X confocal image of endogenously tagged active zone marker Neurexin::Skylan-S in the dorsal nerve cord of an L4-stage worm. Outline of crop from 100X image of the same worm shown in green. Note the lower signal-to-noise ratio compared to B. Scale Bars, 5µm. D. Example Crop from 63X confocal image with annotated ROIs (Green) from different methods showing: *i*. 2D ROIs from Ground Truth annotation, *ii.* Projected 2D ROIs from 1D peak finding algorithm, with valid peaks labeled by black Xs and erroneous peaks (not corresponding to real puncta) labeled with orange Xs, *iii.* 2D ROIs from Global Thresholding algorithm, with missing puncta highlighted in red, and *iv.* 2D ROIs from Local Means Thresholding algorithm, also showing missing puncta highlighted in red. The local means thresholding method demonstrates the highest fidelity to the ground truth annotation. Scale Bar, 5µm. E. Sensitivity (True Positive Rate) for each of the three thresholding methods, measured in the Over Expressed Synaptic Fluorophore Dataset (N=52, red) and the Endogenous Synaptic Fluorophore dataset (N=56, blue). F. Specificity (True Negative Rate) for each of the same three thresholding methods, measured in the Over Expressed Synaptic Fluorophore Dataset (N=52) and Endogenous Synaptic Fluorophore dataset (N=56). G. Balanced Accuracy – the average of Sensitivity and Specificity – for each of the three thresholding methods, measured in the Over Expressed Synaptic Fluorophore Dataset (N=52) and Endogenous Synaptic Fluorophore dataset (N=56).

**Table 1:**
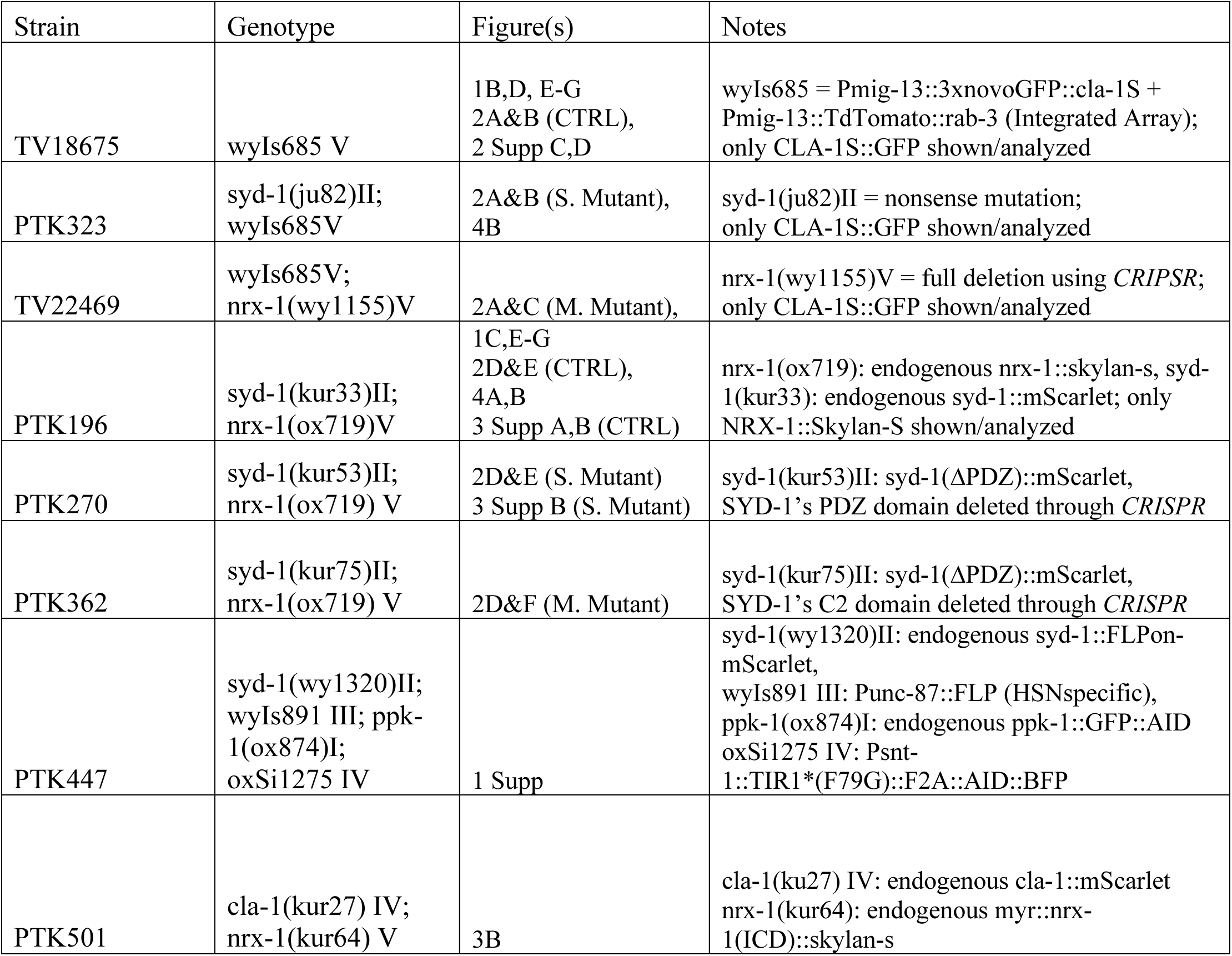
Strain List.

### Software overview

To reduce the barrier to access for unbiased automated puncta quantification in the field, we designed a ‘no-code’ platform for optimizing quantification parameters while also providing a reliable, one-size-fits-all base methodology for quantification. Our software offers a MATLAB-based Graphical User Interface (GUI) tailormade for the detection and characterization of discrete fluorescent puncta in cropped 2D images. The software is distributed both as a MATLAB app and as a stand-alone application, allowing users to run it without requiring a MATLAB license or coding knowledge. Our GUI, *WormSNAP,* implements three main functions: 1) a user-adjustable local means thresholding algorithm for puncta detection, 2) restriction of ROIs based on several characteristics, and 3) visualization of segmented images and associated ROIs. *WormSNAP* enables the detection of puncta from the typical *en passant* synapses (Figure 1) in worm nerve cords, and more complex synapses from outside the nerve cord (Supp Figure 1).

We have also included several ease-of-use features such as the ability to save previously used settings, dataset exclusions, ROI detection parameters, and ROI mask visualizations. Outputs include calculations of commonly quantified parameters such as correlation metrics for 2D images, montages of all analyzed images, summary plots, and organized data tables for downstream analysis using Graphpad Prism or other plotting software. The WormSNAP software, instructions for its usage, and a series of FIJI macros for generating straightened cropped images from a variety of common imaging file formats are all available on a GitHub repository: github.com/Kurshanlab/WormSNAP

## RESULTS

*WormSNAP* has been developed to process cropped and straightened images of up to two fluorescent reporters, hereafter simply termed *crops*. The software was developed using MATLAB R2022a with the Image Processing Toolbox version 11.5 and Statistics and Machine Learning Toolbox version 12.3. The software and associated documentation are available as either a MATLAB add-on or a standalone application on the GitHub repository: github.com/KurshanLab/WormSNAP

In this section, we describe the validation of the thresholding algorithm used by the software and cover the typical pipeline, highlighting useful features such as unbiased ROI curation. We then compare the different 2D correlation metrics available in *WormSNAP* and describe their use cases.

### Thresholding and segmentation algorithm

The first step of ROI detection is thresholding, wherein regions of the image are labeled as foreground or background. Various methods exist for this purpose and can be categorized as either global thresholding, where a single threshold is used for the entire image, or adaptive thresholding, where a different threshold is computed for each pixel in an image. Adaptive thresholds have been shown to be robust to variations in background and low signal-to-noise ratios^13^. Since *crops* from *C. elegans* often exhibit changing backgrounds (due to variations in imaging depth or out-of-focus tissues), sporadic background signals from adjacent regions (e.g. autofluorescent gut granules; Figure 1B,C), and low signal-to-noise ratio especially for endogenous markers^14^(Figure 1C), we predicted that a local means thresholding algorithm would be most appropriate. We chose Bradley’s method, in which a threshold is calculated based on the mean of pixel values in a specified neighborhood around a given pixel^15^. This thresholding algorithm can be adjusted based on the neighborhood size and the threshold sensitivity allowing it to be optimized for different types of fluorescent puncta.

The second step of ROI detection is segmentation, where adjacent puncta that have been grouped into a large ROI are separated into their individual ROIs. Watershed algorithms, which involve the treatment of pixel values as topography, have seen consistent use in image segmentation. The starting points for a watershed algorithm are generally based on the negative of a distance function from the edges of the ROI. We determined that an advanced watershed-based algorithm that uses the Fernand Meyer algorithm and considers both the distance to edge and intensity of a pixel would be most suitable to correctly separate nearby fluorescent puncta^16^.

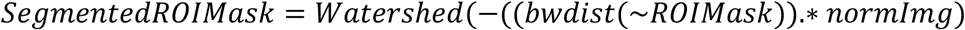

### Method Validation – Comparison of accuracy between thresholding methods

The accuracy of the software was validated on datasets representing two extremes of reporter type: an overexpressed DA9 cell-specific fluorescent protein, mig-13p::CLA-1::GFP, which displays very bright GFP signal, and an endogenously expressed fluorescent protein NRX-1::Skylan-S^17^ which displays dim signal that requires activation from a 405 nm laser in addition to 488 nm excitation to continuously fluoresce. Calculating the ratio of the sum of intensities of pixels assigned (Signal) and not assigned (Noise) to ROIs of the ground-truth annotations for each dataset confirmed that the overexpressed cell-specific reporter exhibits a high signal-to-noise ratio (18.5 ± 1.9) compared to the endogenous reporter (4.09 ± 0.23).

To assess the accuracy of the software compared to the two most prevalent methods in use—1D plot profile analysis (‘peakfinder’) and a global thresholding method—we used receiver operating characteristics, which are based on the ratio of binary classifications of the model to that of the actual classification. These characteristics have been widely used in the assessment of a range of classification methods, such as Information Retrieval, Natural Language Processing, and Machine Learning^18,19^. In our context, they serve to visually represent the likelihood of a method to correctly classify a true signal pixel as part of an ROI (True Positive Rate or TPR) and correctly classify a true noise pixel as background (True Negative Rate or TNR; see methods section).

We first annotated each dataset using a modified version of *WormSNAP* that allows for assisted hand-drawn ROIs, creating the ground truth annotated dataset (Figure 1D*i*). Each dataset was then analyzed using three different thresholding methods. In the first method, a version of *WormSNAP* using a 1D (intensity profile) based detection script was used to identify peaks, and maximal colocalization with the ground truth annotated dataset was determined by varying settings including noise-to-signal ratio, peak prominence, and maximum peak width. Since the 1D plot profile methodology was unable to assign individual pixels as ROIs/nonROIs, we also calculated the metrics in a 1-dimensional manner where a stack of pixels was assigned as ROI (or not) based on the existence of at least one ROI pixel in the stack (Figure 1D*ii*). The same dataset was then analyzed with a version of *WormSNAP* using an Otsu method-based global threshold^20^ (Figure 1D*iii*). Finally, the dataset was analyzed using *WormSNAP* with thresholding settings set to optimize puncta detection in the Control strain for each dataset (Figure 1D*iv*).

The pixel assignment by each method was compared to the hand-annotated ground truth dataset (Figure 1D*i*) to calculate the following metrics: Sensitivity/TPR, Specificity/TNR, and Balanced Accuracy, a metric that combines the Sensitivity and Specificity to assess both aspects of the method.

We compared the *WormSNAP* local means thresholding algorithm to both the 1D peakfinder and the global threshold method in both the overexpressed fluorophore dataset and the endogenous fluorophore dataset (Figure 1E-G). Notably, we observed that while the TNR tends to be high in the 1D peakfinder, it shows significant deficits in TPR, as expected for an overly sensitive method (Figure 1E,F). The global threshold methodology shows the opposite effect, albeit to a much smaller degree. The global threshold is very close to the local means threshold in the overexpressed dataset, demonstrating that it is adequate when dealing with high signal-to-noise images. However, in the low signal-to-noise ratio context of the endogenous reporter, the global threshold displays much lower sensitivity than the local means thresholding, as established in the literature^21^. Thus, the Balanced Accuracy metric, which combines both sensitivity and specificity, suggests that overall, the *WormSNAP* local means thresholding algorithm is superior to either the 1D peakfinder or the global threshold method across a variety of reporter types (Figure 1G).

### Method Validation – Robustness test for local means thresholding

A frequent issue that arises during the development of analysis software for fluorescent imaging is over-optimization, where quantification parameters are overly constrained/fitted to show significant differences between controls and specific mutant phenotypes at the expense of identifying weaker or differing phenotypes^8^. Robustness testing can assess over-optimization by varying ROI detection and selection parameters to determine whether only certain chosen settings result in a particular conclusion.

We evaluated the robustness of WormSNAP by analyzing, for each fluorophore type (overexpressed and endogenous), a dataset consisting of *crops* from multiple genotypes: Control (CTRL) animals, worms containing a mutant allele resulting in a severe phenotype in the fluorescent reporter, and worms containing a mutant allele resulting in a mild reporter phenotype (Figure 2A,E). The strains for each phenotype were chosen based on the extent to which they disrupted fluorescent puncta. The severe phenotype mutant led to defects in protein clustering and thus more diffuse puncta, whereas the mild phenotype mutants only caused reductions in puncta number (compared to controls).

**Figure 2.**
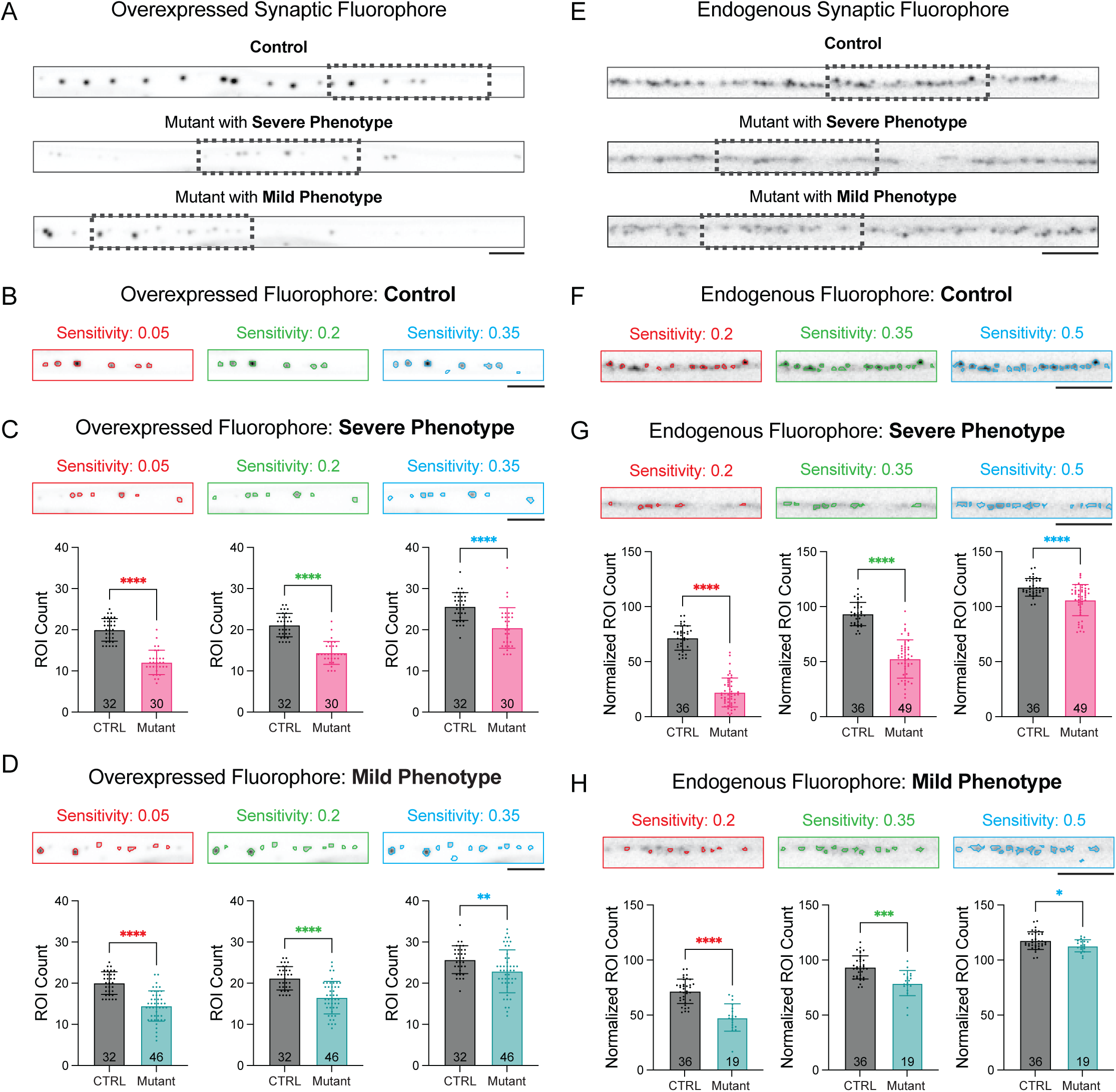
Local Means Thresholding Algorithm shows high robustness. A. Example straightened *crops* of 63X confocal images from the Over Expressed Synaptic Fluorophore Dataset showing CLA-1::GFP-labeled presynapses in the DA9 neurons of the control worms and worms with the different mutant alleles. Scale Bar, 5µm. B. Examples of ROIs detected in the boxed region of the CTRL *crop* (in A) based on minimum (red), ideal (green), and maximum (blue) sensitivity settings for ROI thresholding. Scale Bar, 5µm. C. (Top) Examples of ROIs detected in the boxed region of the Severe Mutant *crop* (from A) based on minimum (red), ideal (green) and maximum (blue) sensitivity settings for ROI thresholding. Scale Bar 5µm. (Bottom) Graphs of ROI counts for CTRL and the severe phenotype mutant at the three sensitivities displayed (*=p<0.05, **=p<0.01, ***=p<0.001, ****=p<0.0001). D. Same as B for the Mild Phenotype Mutant in the Over Expressed Dataset. E. Example straightened *crops* of 100X confocal images from the Endogenous Synaptic Fluorophore Datasets showing NRX-1::Skylan-S-labeled presynapses in the dorsal nerve cords (DNC) of the different mutant alleles. Scale Bar 5µm. F. Examples of ROIs detected in boxed region of the Control *crop* (in E) based on minimum (red), ideal (green) and maximum (blue) sensitivity settings for ROI thresholding. Scale Bar, 5µm. G. (Top) Examples of ROIs detected in the boxed region of the Severe Mutant *crop* (in E) based on minimum (red), ideal (green) and maximum (blue) sensitivity setting for ROI thresholding. Scale Bar, 5µm. (Bottom) Graphs of Normalized ROI count per 100µm (along entire *crop*) for CTRL and mutant with severe phenotype at the three sensitivities picked (*=p<0.05, **=p<0.01, ***=p<0.001, ****=p<0.0001). H. Same as G for the Mild Phenotype Mutant in the Endogenous Dataset.

To test the robustness of our method, we systematically varied each of the following metrics around the original values used for quantification: local means threshold sensitivity, local means neighborhood size, minimum ROI area and maximum length-width ratio. We first optimized the settings for the CTRL *crops* in each dataset, and then, while keeping other parameters fixed, systematically adjusted the parameter of interest around its optimal value. The sensitivity setting was varied over an interval of 0.3 (out of a total possible range of 0-1) in 0.01 increments centered around the optimal sensitivity (0.2 for overexpressed fluorophore and 0.35 for endogenous fluorophore). The ROI counts (or Normalized ROI Counts for Endogenous strains) of each genotype in the dataset was recorded, as well as the p-value derived from a one way anova. We then plotted bar graphs for the minimum, ideal, and maximum sensitivity considered (Figure 2C,D,G,H).

The robustness testing showed that the settings used were not overly sensitive to the comparisons made in this paper for both the overexpressed synaptic fluorophore (Figure 2C, D) and the endogenous synaptic fluorophore (Figure 2G, H). While the individual output values differed notably as the sensitivity was changed, the conclusion of whether the two genotypes were significantly different from each other remained unchanged. In fact, we only saw noteworthy increases in p-value for the Mild Phenotype datasets (Figure 2D,H). In overlaid images of example *crops* using different parameter settings (highlighted in blue in Figure 2B-D and F-H), it appears that the higher sensitivity settings falsely include noise as puncta, thereby inflating the count.

### GUI Pipeline

Upon startup, *WormSNAP* prompts users to select a folder containing preprocessed *crops*. The *WormSNAP* GUI consists of two sections: a display panel and an input panel. The display panel shows *crops* ordered by genotype and number, while the input panel serves as the primary way to interact with the GUI. The input panel contains three tabs that users proceed through sequentially. First, the Preprocessing tab (Supp Figure 3A) allows users to name the image channels, rename genotypes, and, if necessary, specify the magnification/resolution.

**Figure 3.**
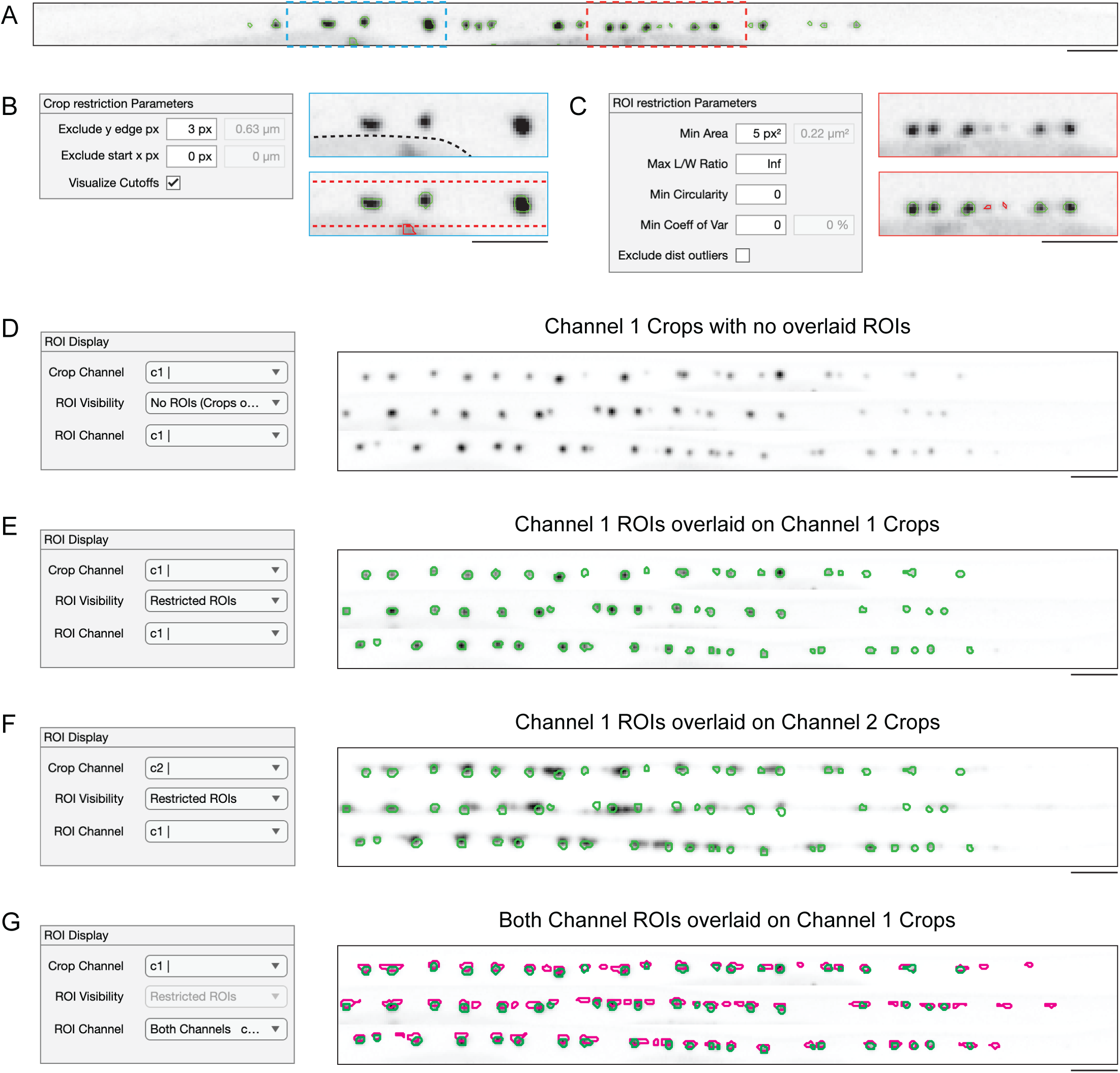
ROI Restriction and Visualization in WormSNAP GUI. B. ROIs detected by Local Area Thresholding algorithm on the *crop* of an Overexpressed Synaptic Fluorophore (previously shown in Figure 1B). C. (Left) Crop Restriction parameters from *WormSNAP’s* ROI Detection Tab (see Figure 4 Supp B), with a y-edge exclusion of 3px (≈0.63µm). (Right Top) Magnified view of the blue box (in A) showing no ROIs assigned and illustrating that the bottom-most punctum is part of the gut (outlined in dotted black line). See Figure 1B for the full image showing the gut impinging on the crop outline. (Right Bottom) Example of how excluding y-edge pixels can remove a misassigned ROI from the gut granule. Dotted lines indicate the boundaries beyond which ROIs are ignored. The excluded ROI is shown in red. Scale Bar, 5µm. D. (Left) ROI Restriction parameters from *WormSNAP’s* ROI Detection Tab, set to a minimum area of 5px^2^/0.22µm^2^. (Right Top) Magnified view of the red box (in A), showing no ROIs assigned. (Right Bottom) Example of how the minimum area setting excludes misassigned ROIs that appear as individual bright pixels due to noise. Excluded ROIs are shown in red. Scale Bar, 5µm. E. Magnified view of *WormSNAP’s* Results Tab (see Figure 4 Supp C) and an example 2-channel *crops* visualization from the Display Tab, showing Channel 1 (CLA-1::GFP) intensities for 3 *crops*. Scale Bar, 5µm F. Magnified view of *WormSNAP’s* Results Tab (see Figure 4 Supp C) and an example 2-channel *crops* visualization from the Display Tab, with (green) ROIs of Channel 1 (CLA-1::GFP) overlaid on Channel 1 (CLA-1::GFP) *crops*. Scale Bar, 5µm G. Magnified view of *WormSNAP’s* Results Tab (see Figure 4 Supp C) and an example 2-channel *crops* visualization from the Display Tab, with the (green) ROIs of Channel 1 (CLA-1::GFP) overlaid on Channel 2 (RAB-3::tdTomato) *crops*. Scale Bar, 5µm H. Magnified view of *WormSNAP’s* Results Tab and example 2-channel *crops* visualization from the Display Tab, showing ROIs from both channels overlaid on the Channel 1 (CLA-1::GFP) *crops*: Channel 1 (CLA-1::GFP) ROIs in green and Channel 2 (RAB-3::tdTomato) ROIs in magenta. Scale Bar, 5µm

Users then proceed to the ROI detection tab (Supp Figure 3B) to specify the thresholding and restriction settings for each channel. The restriction parameters help exclude spurious ROIs that arise from imaging artifacts or noise (Figure 4A). The first set of restriction parameters, called Crop Restriction Parameters, enable the exclusion of ROIs based on their proximity to the edges of *crops*, which helps exclude artifacts such as *C. elegans* autofluorescent gut granules (Figure 3B). The second set of parameters filters ROIs based on features including area, circularity, length-to-width ratio, and intra-ROI intensity variance. These criteria allow users to exclude noise that often appears as a few bright pixels (Figure 3C).

**Figure 4.**
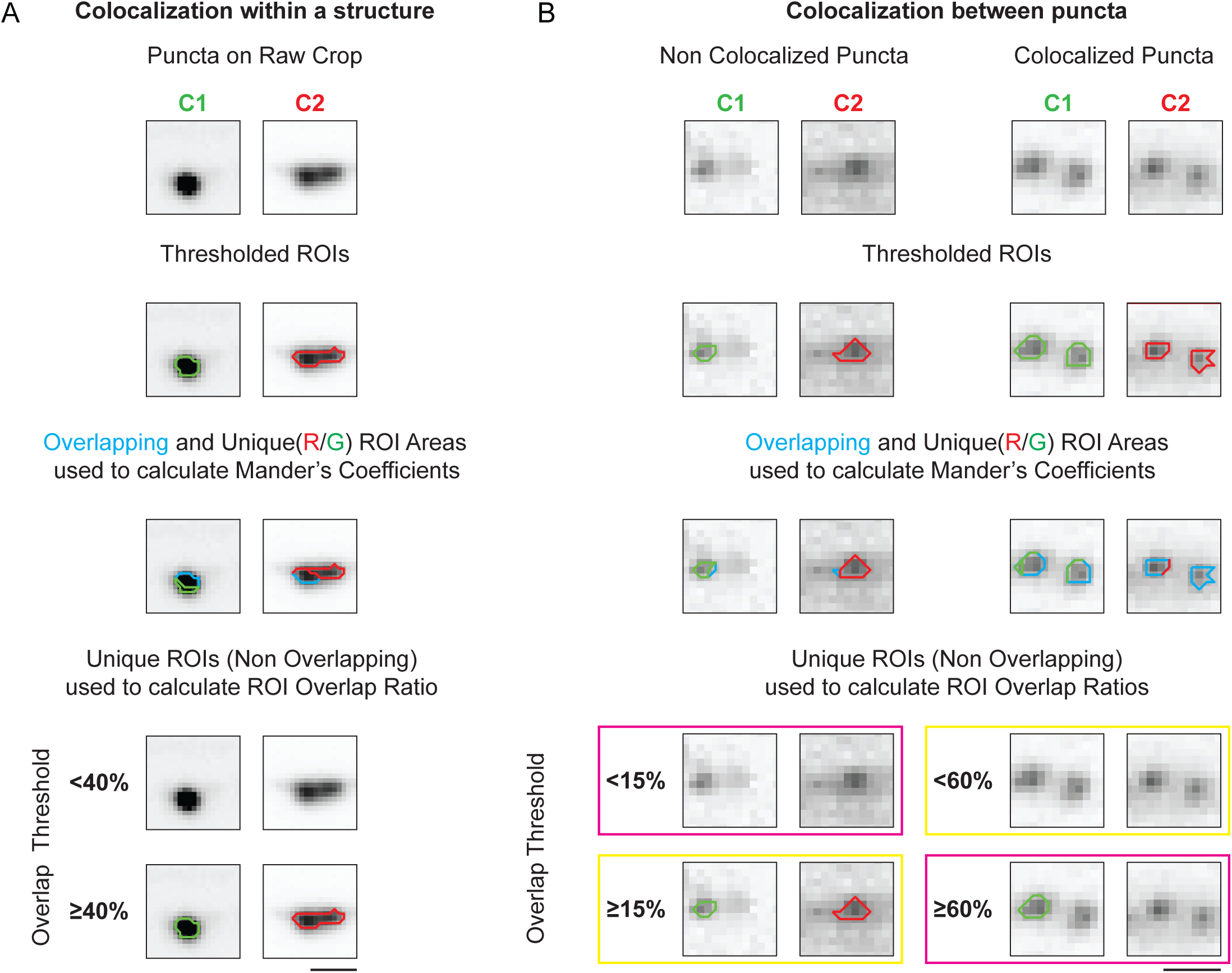
Comparison of WormSNAP Correlation Metrics. A. Example of two proteins (CLA-1, an active zone marker, and RAB-3, a vesicle marker) that exhibit different localization patterns *within* the same synaptic structure. Mander’s coefficients (the ratio of overlapping area/total ROI area in each channel) accurately capture these differences because the extent of overlap between the channels is markedly different. In contrast, the ROI Overlap Ratio, which provides binary information for individual puncta, either shows complete colocalization at Overlap Thresholds (OT) below 40%, or none at Ots of 40% or more. Therefore, for assessment of colocalization *within* puncta (or structures), Mander’s coefficient are more suitable. Scale Bar, 2µm. B. Example of two proteins (CLA-1 and NRX-1) that exhibit different localization patterns across synapses in a mutant background – i.e., they are no longer found at the same synapses. Two scenarios are shown: non-overlapping puncta (left) and overlapping puncta (right). Mander’s Coefficients yield a non-zero value for the non-overlapping puncta, owing to minor boundary overlap, and a value below 1 (i.e., less than complete colocalization) for the overlapping puncta. In contrast, the ROI Overlap Metrics use an Overlap Threshold (OT) to determine colocalization. Correct assignments by the ROI Overlap Metrics are indicated by the yellow rectangle; incorrect assignments are pink. Non-overlapping puncta are correctly identified at OTs of 15% or higher, while overlapping puncta are correctly identified below an OT of 60% - indicating that the optimal OT for this dataset lies between 15% to 60%. Scale Bar, 1µm.

After ROIs are detected in all available channels, users proceed to the Results tab (Supp Figure 3C) to visualize the ROIs and configure output settings. Using the ROI Display panel, users can easily view ROIs. When viewing 2-channel images, users can view ROIs of each channel (Figure 3C) as well as mix and match between ROIs and Channels, even allowing users to view ROIs from both channels overlaid on top of a crop (Figure 3D-G).

*WormSNAP’s* output options (Supp Figure 3C) include .csv files of selected quantification parameters, a combined .csv file containing ROI-specific data for further analysis, a ‘.mat’ file that allows you to easily replicate previous analyses, and PDF-ready files of montages or individual *crops* with ROI overlays of the user’s choice (Supp Figure 3F). A step-by-step walkthrough of the software is available in the GitHub documentation.

### GUI Validation – Scalability

Scalability of a method is important for widespread adoption. We optimized the *WormSNAP* GUI to analyze datasets containing more than 100 *crops*. One of the GUI’s main advantages is the ability to easily view many *crops* at once; however, this can be computationally intensive, leading to software slowdowns or crashes for large datasets. To address this, we implemented a tab system to limit the maximum number of *crops* displayed simultaneously. We compared a single-tab display of all *crops* to a split-tab display (with 50 crops per tab) for datasets 50 to 200 *crop*s (Supp Figure 3D, E). Both the time-to-display and ROI calculation time decreased when using split tabs instead of a single tab as the total number of *crops* increased, demonstrating that the split tab method is more effective for displaying large datasets.

### Colocalization metrics

In fluorescence microscopy, assessing colocalization between two fluorescent signals is crucial for understanding biological interactions of the underlying proteins^22^. Hence, WormSNAP also calculates correlation metrics for 2D images as part of its output (Supp Figure 3C,F). Two metrics widely used in the field are the Pearson Correlation Coefficient (PCC) and the Manders’ Coefficients, M1 and M2^23^.

The PCC quantifies the linear relationship between the intensities of two channels, ranging from -1 to 1, where values close to 1 indicate a strong positive correlation^24^. It accounts for the pixel intensities of each channel over the entire image. In unprocessed images, the PCC can be severely impacted by ordered background fluorescence - such as that caused by the sample’s 3D geometry - as well as by noise^25^. However, because our workflow preprocesses images into *crops* that minimize background area, we can substantially reduce the impact of noise and unrelated structured background (Figure Supp 4A,B). For our crop images, the main limitation of PCC lies in its single-metric output encompassing both channels, rendering it incapable of distinguishing which channel exhibits disruption.

In contrast, Manders’ Coefficients measure the fraction of signal in one channel that overlaps with the signal in the other, with M1 quantifying the fraction of signal pixels in channel 1 that are also signal pixels in channel 2, and M2 measuring overlap in the opposite direction^26^. Because M1 and M2 provide channel-specific information, they are well suited to robust thresholding algorithms such as the local means algorithm used by *WormSNAP*.

*WormSNAP* also calculates a novel third set of metrics, the ROI Overlap Ratios, R1 and R2. R1 is the fraction of ROIs (as opposed to pixels) in channel 1 that overlap with ROIs in channel 2, above a user-defined Overlap Threshold. The Overlap Threshold defines the minimum pixel overlap needed to classify two ROIs as overlapping; a lower threshold makes partially overlapping markers more likely to be considered co-localized. R2 measures the overlap in the opposite direction.

To assess changes in the colocalization pattern between two proteins that occupy different regions *within* the same structure, such as active zone proteins (e.g. CLA-1) localized to a distinct region within the synaptic bouton, and synaptic vesicle markers (e.g. RAB-3) that fill the bouton, the Manders’ Coefficients are more suitable. This is because they quantify the *relative* overlap between ROIs (Figure 4A), allowing researchers to determine whether this balance is altered in a particular mutant.

To determine whether two proteins are located within the same structure – for example, whether different synapses contain one protein versus another – the ROI Overlap Ratios are preferred. This is because they assign individual, binary colocalization values for individual ROIs and disregard small overlaps between adjacent but non-overlapping ROIs in different channels (Figure 4B). The ROI Overlap Ratios can be further tuned for specific pairs of markers by setting an Overlap Threshold that distinguishes non-overlapping puncta as separate while still classifying truly overlapping puncta together (in Figure 4B, this threshold ranged from 15% -60%).

Noise is a major consideration when selecting a correlation metric^22^. We generated synthetic two-channel datasets based on Channel 1(CLA-1::GFP) of the Overexpressed Synaptic Fluorophore CTRL *crops* with four different noise types designed to mimic typical confocal microscopy issues: Thermal noise, Shot noise, Salt and Pepper noise, and Speckle noise (Supp Figure 3C). The PCC, M1, and R1 were then computed to measure the correlation between the original image and the noisy image. Because the PCC ranges from -1 to 1 – twice the range of M1 and R1 (which span [0,1]) – it was normalized to [0,1]. Next, the metrics were plotted for the different correlation coefficients to determine which metric remained closest to perfect correlation (=1) under various noisy conditions. M1 performed the worst under all noise types except salt and pepper, where PCC fared worse (Supp Figure 4D). PCC and R1 showed the least deviation overall, with R1 remaining unaffected by salt and pepper noise.

In summary, *WormSNAP* provides a range of colocalization metrics that balance accuracy and noise sensitivity, enabling selection of the most suitable measurement for each dataset and scientific question.

## DISCUSSION

In this paper, we introduce *WormSNAP*, an open-source graphical user interface (GUI) for no-code, local means-based region of interest (ROI) detection, optimized for analysis of large datasets of fluorescent puncta in *C. elegans*. We highlight the strong performance of our local means thresholding algorithm relative to existing methods, especially across a range of fluorescent markers with varying signal-to-noise ratios. To our knowledge, this is the first GUI that allows simultaneous display and interaction with images and 2D ROIs from entire datasets. We have also incorporated multiple user-friendly features such as saving thresholding parameters, renaming channels, managing outliers, selecting correlation metrics, exporting data for easy plotting, and saving images in figure-friendly formats. Through these advancements, we aim to standardize puncta detection in *C. elegans* research and substantially shorten the time from image acquisition to publishable data, ultimately accelerating research workflows.

### Interpretation of results

By comparing model accuracy with other methods, we demonstrate that WormSNAP’s local means thresholding algorithm is well-suited for puncta detection and characterization when compared to the most used methods. In addition, our results show that WormSNAP’s underlying methodology is highly robust: user-modifiable settings support the quantification of a wide range of fluorophores imaged in *C. elegans* nerve cords and more complex synapses.

### Advantages and limitations of the software

Our software offers direct control over ROI detection and restriction parameters without coding requirements, as well as the ability to analyze large datasets. Thus, users can easily optimize the puncta detection settings while avoiding the barriers of specialized programming knowledge. Additionally, our methodology exhibits consistent improvement over widely used approaches, such as 1D plot profiles and global threshold-based ROI detection. With various ease-of-use features, users can annotate datasets and obtain useful outputs without recalculating data or learning how to process esoteric file types.

Currently, WormSNAP is limited to 2D puncta detection of up to 2 channels at a time, constraining its applicability to certain multicolor imaging strategies^27^. Also, the preprocessing step of drawing *crops* precludes a fully automated pipeline. However, time spent analyzing images is still significantly reduced, because *crops* need to be drawn only once per dataset. Analysis parameters and exclusions are also saved after the session is done, allowing for re-analysis of older data using new optimized parameters.

Although other software (e.g. *WormPsyQi* ^28^) can detect puncta in 3D, it requires overexpressed cytoplasmic markers to define the axon boundary, and the synapse detection model may require further training to function. In contrast, *WormSNAP* was designed as an easy-to-use software that can be applied to existing fluorescent imaging datasets without axonal boundary demarcation. Moreover, *WormSNAP* lets users display large datasets effortlessly and repeat the same analyses with a single click.

### Future directions and potential improvements

Future work will focus on extending our approach to 3D images. Currently, maximum intensity projection from 3D image stacks is used, but for *C. elegans* nerve cords (∼40 nm thick), the need for 3D data is relatively low^29^. Ongoing efforts aim to enable 3D thresholding in future releases^25^. Additional improvements under development include blinding functions and automated puncta detection in standard confocal images using feature detection methods.

## METHODS

### C. elegans methods

*C. elegans* strains were derived from the Bristol strain N2 and raised at 23°C on NGM plates seeded with OP50 *E. coli*, according to standard protocols^30^. The ‘Overexpressed Cell Specific Fluorescent Protein’ datasets were collected by imaging CLA-1::GFP (a presynaptic active zone marker^4^) in the axon of the DA9 neuron from the following strains: TV18675 [*wyIs685 V*], an integrated transgene that expresses CLA-1::GFP driven by the DA9 neuron-specific mig-13 promoter; TV22469 [*wyIs685 V; nrx-1(wy1155) V*] (‘Mild Phenotype’), which expresses the same transgene in a *nrx-1(-)* background showing loss of posterior axonal CLA-1::GFP punta^12^; and PTK323 [*wyIs685 V;syd-1(ju82) II*]^12^ (‘Severe Phenotype’), which expresses the same transgene in a *syd-1(-)* background, leading to loss of most CLA-1::GFP puncta.

The ‘Endogenous Fluorescent Protein’ dataset was collected by imaging endogenously-tagged NRX-1::Skylan-S in the dorsal nerve cord of worms from the following strains: PTK196 [*syd-1(kur33) II ;nrx-1(ox719) V*] (‘Control’), which expresses endogenously tagged NRX-1::Skylan-S and endogenously tagged SYD-1::mScarlet; PTK362 [*syd-1(kur75) II; nrx-1(ox719) V*] (‘Mild Phenotype’), which expresses endogenously tagged NRX-1::Skylan-S and endogenously tagged SYD-1(ΔC2)::mScarlet and exhibits reduced number and intensity of NRX-1::Skylan-S puncta; and strain PTK270 [*syd-1(kur57)II ;nrx-1(ox719)V*] (‘Severe Phenotype’), which expresses endogenously tagged NRX-1::Skylan-S and endogenously tagged SYD-1(ΔPDZ)::mScarlet and exhibits a severe disruption of NRX-1::Skylan-S clustering.

The dataset used in Figure 4B was collected by imaging NRX-1::Skylan-S and CLA-1::mScarlet in the dorsal nerve cord of worms from strain PTK501 [cla-1(kur27) IV; nrx-1(kur64) V], which expresses endogenously tagged CLA-1::mScarlet and Myristoylated::NRX-1(Intracellular Domain)::Skylan-S in the NRX-1 locus and exhibits loss of synaptic (CLA-1 colocalized) NRX-1 puncta.

Images of non-nerve cord synaptic puncta were collected from PTK447 [*syd-1(wy1320) II; wyIs891 III; ppk-1(ox874); oxSi1275 IV*], which expresses SYD-1::FLPon-mScarlet in the HSN neuron with the HSN-specific unc-86 promoter driving FLP recombinase.

### Data acquisition and crop tracing

Confocal images of fluorescently labeled proteins were collected at room temperature in living, anesthetized *C. elegans.* L4.4-stage^31^ hermaphrodites were anesthetized in 20 mM levamisole (Sigma-Aldrich) in M9 buffer, mounted on 10% agarose pads, and sealed under a #1.5 coverslip with petroleum jelly.

Images were acquired on a spinning disk confocal (3i) equipped with an Evolve 512 EMCCD camera (Photometrics) and a CSU-X1 M1 spinning disk (Yokogawa) mounted on an inverted Zeiss Axio Observer Z1 microscope. The ‘Overexpressed Cell Specific Fluorescent Protein’ datasets were collected with a Zeiss Plan-Apochromat 63x/1.4NA oil immersion objective in z-stacks of the DA9 axon in the dorsal nerve cord, with 0.27 μm z steps over a 5 μm z range. The ‘Endogenous Fluorescent Protein’ images were collected with a Zeiss Plan-Apochromat 100x/1.4NA oil immersion objective in z-stacks from the dorsal nerve cord, with 0.27 μm z steps over a 6 μm z range. Z-stacks were obtained centered around the highest in-frame fluorescence of either the DA9 axon or DNC using non-saturating laser, exposure and camera settings. Synapses in the HSN neuron were collected in z-stacks with 0.27 μm z-steps over an 8.4 μm z range, also with a 100x/1.4NA oil immersion objective. Images were acquired using Slidebook 6.0 software (3i).

During initial image processing procedures, raw z-stack volumes from the native .sld Slidebook file format (3i) were rendered into 16-bit maximum intensity projections tifs, and 20-pixel-wide lines were drawn over synaptic regions of interest and straightened into 32-bit tif cropped images using Fiji^32^ macro ‘Batch Axon Tracer - sld’ included in the Github repository.

### Method Validation – Robustness test for local means thresholding

Using a custom ROI overlap analysis script, the different methods were compared *crop*- wise to the ground truth annotated dataset, by evaluating three standard receiver operating characteristics for accuracy comparisons, namely the Sensitivity/True Positive Rate (TPR), Specificity/True Negative Rate (TNR) and Balanced Accuracy (BA).

The receiver operating characteristics were calculated as follows: Sensitivity/True Positive Rate (TPR) was assessed as:

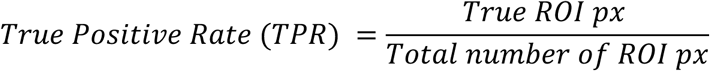

where ‘True ROI px’ refers to an ROI pixel that colocalized with a ground-truth annotated ROI pixel,

Specificity/True Negative Rate (TNR) was assessed as:

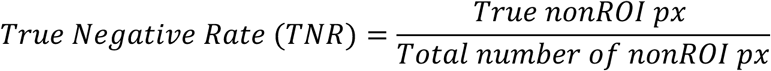

where ‘True nonROI px’ refers to a pixel that was not considered as part of an ROI by either the software or the ground-truth annotation.

Balanced Accuracy (BA) was assessed as follows:

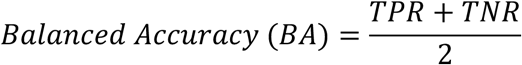

### 2D channel correlation analysis – Correlation metrics

*WormSNAP* calculates three different correlation metrics: the Pearson’s Correlation Coefficient, the Manders’ Coefficients and the ROI Overlap Ratios.

The Pearson’s Correlation Coefficient is calculated directly from ROI intensity as follows for a two-channel image:

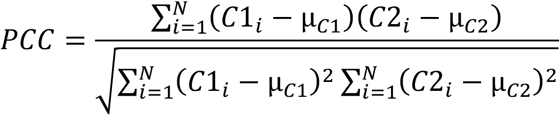

Where:

N is the total number of pixels in the image
*C*1*_i_* is the intensity of pixel *i* in Channel 1
µ*_c_*_1_is the mean intensity in Channel 1
*C*2*_i_* is the intensity of pixel *i* in Channel 2
µ*_c_*_2_is the mean intensity in Channel 2

The Manders’ Coefficients and ROI Overlap Ratios are calculated based on the ROI assignments post thresholding and ROI restriction.

The Mander’s Coefficients (M1 and M2) are calculated as follows for a two-channel image^26^:

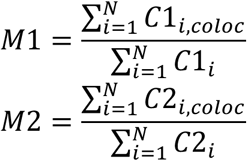

Where:

is the total number of pixels in the image
*C*1*_i_* = 1 if pixel *i* is thresholded as signal in Channel 1 and *C*1*_i_* = 0 otherwise
*C*2*_!_* = 1 if pixel *i* is thresholded as signal in Channel 1 and *C*2*_i_* = 0 otherwise
*C*1*_i,coloc_* = 1 if pixel *i* is thresholded as signal in both channels and *C*1*_i,coloc_* = 0
therwise
*C*2*_i,coloc_*= 1 if pixel *i* is thresholded as signal in both channels and *C*2*_i,coloc_* = 0
therwise

The ROI Overlap Ratios (R1 and R2) are calculated as follows for a two-channel image:

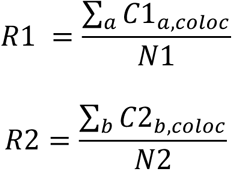

Where:

N1 is the number of ROIs detected in channel 1 N2 is the number of ROIs detected in channel 1
*C*1*_a,coloc_* = 1 if ROI *a* is considered to have significant overlap with ROIs in Channel 2 based on an Overlap Threshold of the percentage of its pixels thresholded as signal in Channel 2, *C*1*_a,coloc_* = 0 otherwise
*C*2*_b,coloc_* = 1 if ROI *b* is considered to have significant overlap with ROIs in Channel 1 based on an Overlap Threshold of the percentage of its pixels thresholded as signal in Channel 1, *C*2*_b,coloc_* = 0 otherwise

### 2D channel correlation analysis – Synthetic noise

Noise was added to original images using Matlab’s Image Processing Toolbox’s *imnoise* function. The function had inbuilt settings for Salt and Pepper noise and Speckle noise while Thermal noise was modeled using a Gaussian distribution and Shot noise was modeled using a Poisson distribution^33^. The default settings were used for the Poisson and Speckle Noise. The Gaussian function used had mean of 0 and variance of 0.01 while the Salt and Pepper Noise had a density of 0.01.

### Statistical Analysis

All statistical analysis was done using Prism. For Figure 2, Brown-Forsythe and Welch One Way Anova tests were carried out at. The Dunnett’s T3 multiple comparisons test was then done between the CTRL and each mutant phenotype strain and the resulting p-value plotted as appropriate. For Supplement Figure 4, a paired two-tailed Student’s *t test* was used to calculate the p-values.

---

**Figure 1 Supplement:**
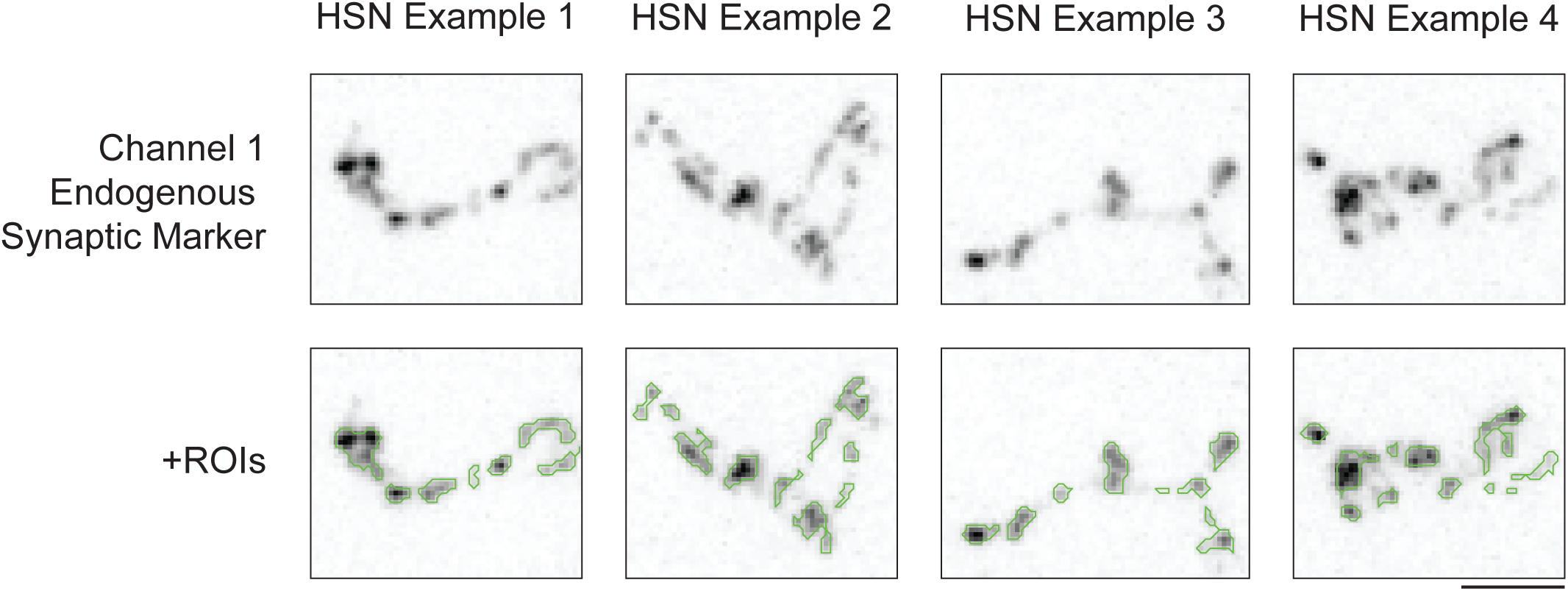
Local Means Thresholding analysis of ROIs from HSN synapses. Example *crops* from HSN synapse from worms with endogenous expression of a synaptic fluorophore (SYD-1::mScarlet). Top: *Crops*. Bottom: *Crops* with ROI overlay (green). Scale, 3 µm.

**Figure 3 Supplement:**
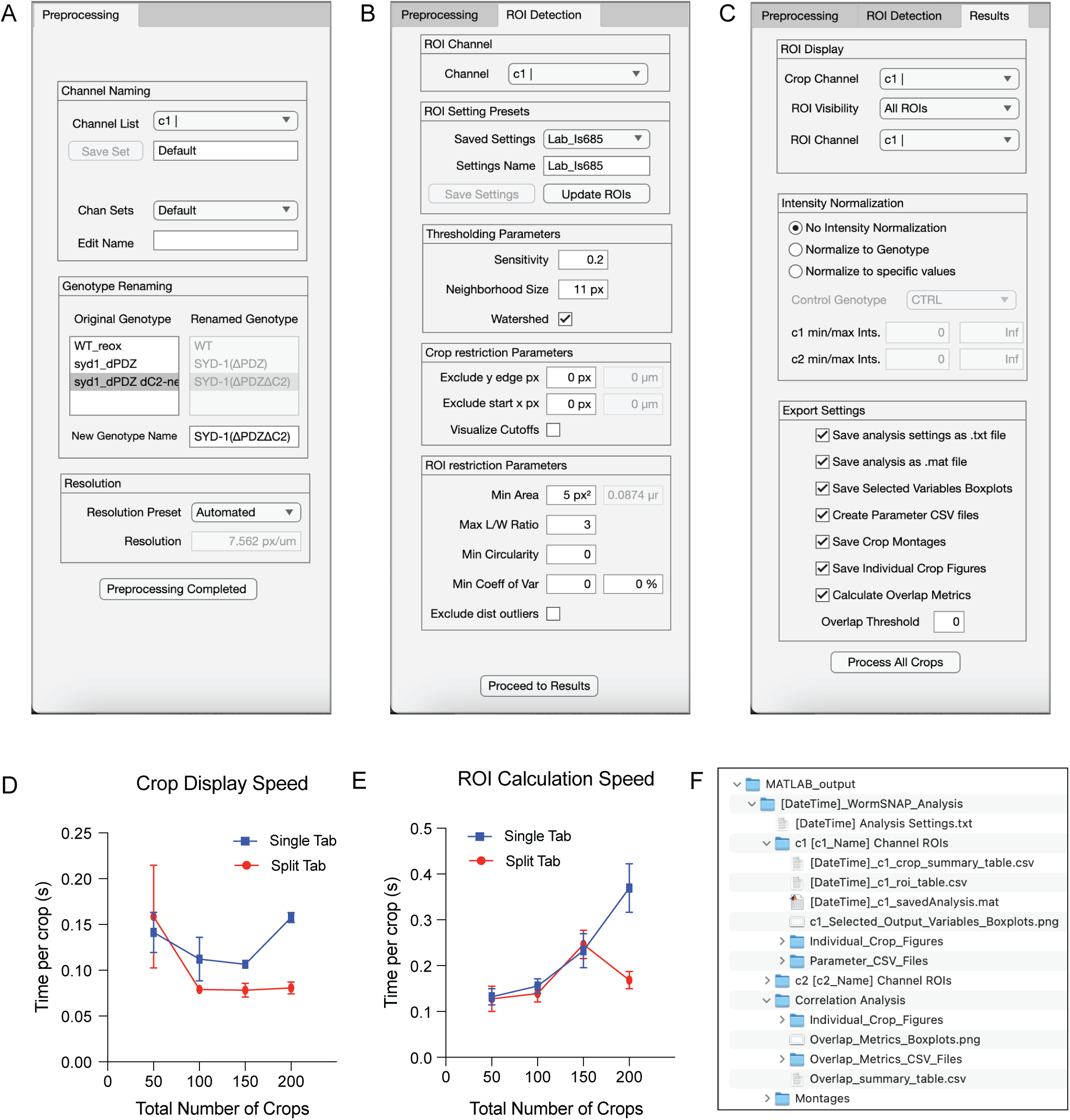
WormSNAP GUI Display and Outputs. A. Screenshot of the Preprocessing Tab in the Graphic User Interface (GUI). B. Screenshot of the ROI Detection Tab in the GUI. C. Screenshot of the Results Tab in the GUI. D. *Crop* Display Speed, measured in seconds per *crop,* for datasets of various sizes displayed in a single-tab versus a split-tab. The split-tab approach limits each tab to a maximum of 50. E. ROI Calculation speed (seconds per *crop)* for datasets of various sizes displayed in a single-tab versus a split-tab. F. Example Output folder for a 2-channel dataset generated using the output settings shown in (C)

**Figure 4 Supplement:**
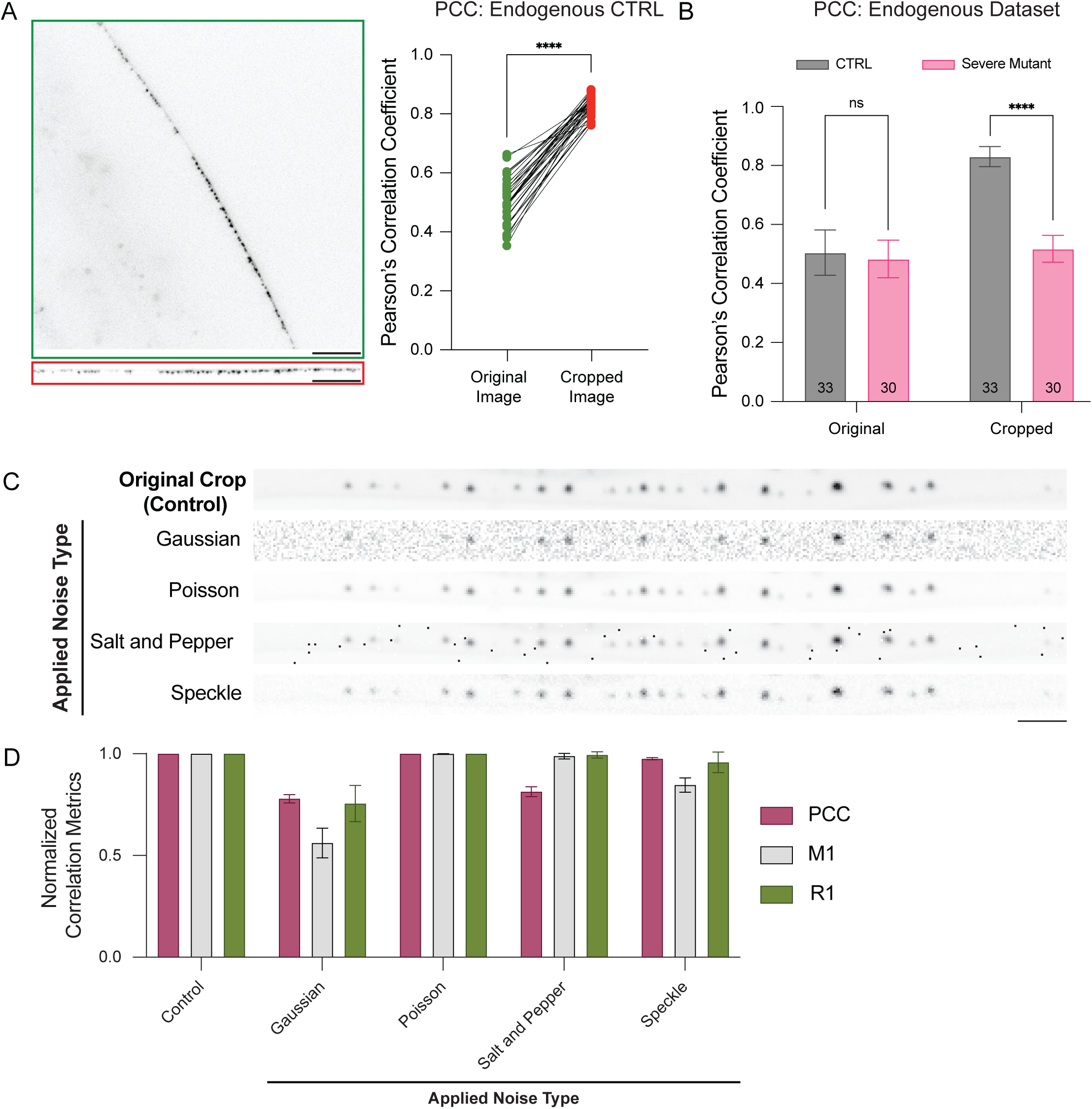
Effects of cropping and noise on WormSNAP Correlation Metrics. A. (Left) Pearson’s Correlation Coefficient (PCC) for original versus cropped images from the endogenous dataset (CTRL). (Right) Example image with the corresponding crop. Scale Bars, 5µm. B. PCC for original and cropped images of the Endogenous dataset, demonstrating that cropping images improves the PCC metric in severe mutant phenotypes but not in CTRL. Consequently, *crops* show a significant difference in PCC, whereas original images do not (*=p<0.05, **=p<0.01, ***=p<0.001, ****=p<0.0001). C. Example images illustrating how different noise types affect the signal from the *crops*. Scale Bar, 5µm. D. Quantification of three Correlation Metrics – PCC, Manders’ Coefficient (M1), and ROI Overlap ratio (R1) – for the various types of noise shown in (C), applied to Channel 1 of an Over Expressed Fluorophore CTRL Dataset. PCC was normalized to [0,1] from [-1,1] for the graph.

